# Multi-season analysis reveals hundreds of drought-responsive genes in sorghum

**DOI:** 10.1101/2025.06.27.662006

**Authors:** Benjamin Cole, Wenxin Zhang, Junming Shi, Hao Wang, Christopher Baker, Nelle Varoquaux, Joy Hollingsworth, Robert Hutmacher, Jeffery Dahlberg, Grady Pierroz, Kerrie W. Barry, Vasanth Singan, Yuko Yoshinaga, Christopher Daum, Matthew Zane, Matthew Blow, Ronan O’Malley, Shengqiang Shu, Jerry W. Jenkins, John T. Lovell, Jeremy Schmutz, John W. Taylor, Devin Coleman-Derr, Axel Visel, Peggy G. Lemaux, Elizabeth Purdom, John P. Vogel

## Abstract

Persistent drought affects global crop production and is becoming more severe in many parts of the world in recent decades. Deciphering how plants respond to drought will facilitate the development of flexible mitigation strategies. *Sorghum bicolor* L. Moench (sorghum), a major cereal crop and an emerging bioenergy crop, exhibits remarkable resilience to drought. To better understand the molecular traits that underlie sorghum’s remarkable drought tolerance, we undertook a large-scale sorghum gene expression profiling effort, totaling nearly 1,500 transcriptome profiles, across a 3-year field study with replicated plots in California’s Central Valley. This study included time-resolved gene expression data from roots and leaves of two sorghum genotypes, BTx642 and RTx430, with different pre-flowering and post-flowering drought-tolerance adaptations under control and drought conditions. Quantification of genotype-specific drought tolerance effects was enabled by *de novo* sequencing, assembly, and annotation of both BTx642 and RTx430 genomes. These reference-quality genomes were used to construct a pan-gene set for characterizing conserved and genotype-specific expression. By integrating time-resolved transcriptomic responses to drought in the field across three consecutive years, we identified a set of drought-responsive genes that responded similarly in all three years of our field study. This expansive dataset represents a unique resource for sorghum and drought research communities and provides a methodological framework for the integration of multi-faceted time-resolved transcriptomic datasets.

## Introduction

To meet the ever-growing global demand for resources (including food, feed and energy), there is an urgent need to explore alternative and renewable feedstocks while limiting the negative impacts of their production [1]. Bioenergy is an attractive source of renewable fuels [2], with bioethanol production already ubiquitous in the energy market. However, biomass production costs, yields, and conversion efficiencies still need to be substantially improved for cellulosic-derived bioenergy to be a viable alternative to current, dominant energy sources. This improvement would ideally incorporate crop resilience under extreme growing conditions. Drought, for example, poses especially large challenges to biomass crop production, resulting in up to 24-27% losses in yield [3]. Sorghum [*Sorghum bicolor* (L.) Moench] is an important cereal crop for much of the developing world, a large component of animal feed for developed nations, and a promising bioenergy resource [4]. This is in part due to its high resilience to poor growing conditions, including drought and marginal soils, as well as its ease of cultivation [5]. Its relatively high yield under drought conditions is of particular interest, as droughts have become more frequent and more intense recently in many parts of the world [6]. Despite the ability of sorghum to persist through drought imposed at various life stages (*e.g.,* pre-flowering and post-flowering), severe drought in most cases reduces grain production and overall biomass growth [7]. Taken together, these features, combined with the relatively small, diploid sorghum genome, large germplasm collections, and efficient transformation and editing methods [8,9], make sorghum an excellent model for studying drought tolerance in agricultural systems.

Recently, a large consortium set out to profile genome-wide responses to sorghum plants experiencing drought using advanced ‘omics technologies [10–22]. For this project sorghum plants were grown in replicated agricultural plots located at the Kearney Agricultural Research and Extension Center of the University of California’s Division of Agriculture and Natural Resources (UC-ANR) in Parlier, CA. This location was chosen due to its well-drained sandy-loam soil, high summer temperatures that reflect future climatic conditions and, most importantly, the near-total lack of precipitation during the summer months that make it possible to tightly control the amount of water applied. Transcriptomic data from a single year [11] of this experiment revealed that close to 30% of the whole transcriptome was altered by drought in two sorghum cultivars: BTx642, a stay-green [23] variety that retains photosynthetic potential during severe post-flowering drought stress, and RTx430 that lacks this ability. One of the notable advantages of this study included conducting the field study under “real” drought conditions, which greenhouse or in vitro settings simply cannot reproduce [24]. In addition to leaves, this study also concurrently sampled the biotic environment of sorghum roots (the rhizosphere), the microorganisms that live in the soil near plant roots and interact with the plant and react to drought conditions [13,16]. While highly informative, this study did not consider year-to-year variability in weather and soil conditions, which can be large and undoubtedly impact drought severity, microbial community composition, as well as confounding environmental stresses (e.g., heat). Sampling across multiple growing seasons and/or in different locations can provide a more robust evaluation of plant performance during drought.

Moreover, in the first report transcriptomic data was only mapped to the BTx623 reference genome [25,26]. Using a single reference genome for transcriptional analysis of different genotypes introduces several potential artifacts. First, pan-genomic analysis of several plants has revealed extensive intraspecific presence-absence variation [27]. Thus, using a reference genome from a different genotype will completely miss genes that are not found in both genotypes. Second, genes in their degree of DNA similarity, which will affect measured expression levels due to the accuracy of read mapping. Finally, copy number variation will introduce errors into expression measurements. While the overall errors due to these factors may be relatively small on a genome-wide basis, they will be exaggerated for divergent genes which in some cases may be the most important genes determining genotype-specific phenotypes. Thus, a pan-genomic approach is necessary to fully understand the molecular shifts that occur during drought across multiple genotypes, even if they are closely related [28].

Here, we incorporate two additional years of data into the transcriptome analysis of field-droughted sorghum [11] and use a comparative genomics approach to more accurately characterize gene expression in each of the genotypes used. We also introduce additional time points around the water transitions (i.e., turning water back on after pre-flowering drought, and turning water off during post-flowering drought) to capture rapid responses following the end or onset of drought. Using this data, we defined a robust and comprehensive transcriptomic resource that includes highlighting loci that are highly sensitive to drought.

## Results

### Drought treatments and sampling

As results from field studies strongly depend on variable soil and weather conditions, we decided to add two additional seasons of sampling on top of the initial season previously reported[11] to complete a 3-year drought trial. The present study specifically included sampling of two tissue types (root and leaf), two genotypes (RTx430 and BTx642), and three watering regimes (well-watered, pre-flowering drought and post-flowering drought), with weekly sampling across 17 weeks in a matrixed experimental design conducted on at UC-ANR Kearney Agricultural Research and Extension Center plots in the California Central Valley (Parlier, CA) (**Fig. 1**). For both types of drought, the transition in watering status occurred 8 weeks after planting (Day 57), with pre-flowering drought plots receiving a full water treatment for the first time 7 weeks after planting, while regular watering ceased for post-flowering droughted plots at that time. Compared to the first year’s sampling procedure, we increased sampling density in Years 2 and 3, above the previous once-per-week regime, around the drought transition (8 hours, 1 and 3 days after watering resumption for pre-flowering plots, 2, 5 days after stopping watering for post-flowering plots). This allowed us to gain finer-scale resolution of transcriptomic changes after the resumption of watering after pre-flowering drought (Pre-flowering Drought Recovery) or after the onset of post-flowering drought. In total, we employed RNA-seq to characterize the transcriptomes from 1,476 samples.

**Figure 1:**
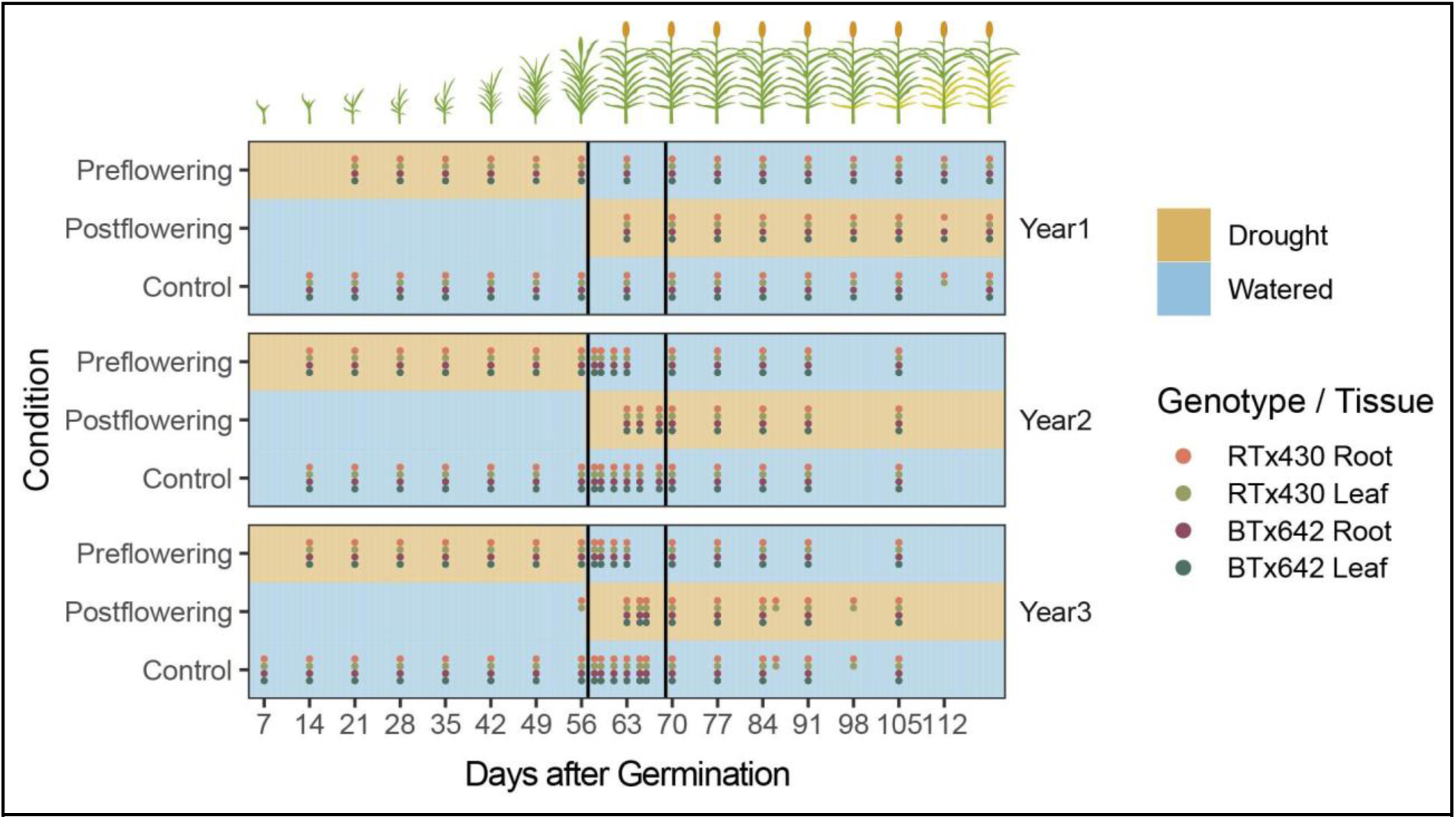
Experimental Overview. Samples were taken weekly each year, starting 7 days after germination. Blue regions in this diagram indicate periods of watering for each condition, while brown regions indicate periods of drought (no watering). Denser sampling occurred around the watering (and flowering) transition. Dots are colored according to the tissue and genotype sampled for a given week.

### A comparative genomic approach for transcriptome analysis

While using a single genomic reference for a multi-genotype study simplifies the analysis of gene expression variation across multiple genotypes by ignoring gene presence/absence and copy number variation, genotypic differences important for the array of phenotypes observed, especially during drought (*e.g.,* stay-green), that are encoded by genotype-specific alleles, or copy-number variation would be missed. To address these shortcomings, we generated *de novo* genome assemblies and annotations for both BTx642 and RTx430 genotypes using a combination of shotgun DNA sequence data, HiC data, RNA-seq data from the 3-year EPICON time course survey (current study), as well as Iso-seq data derived from total RNA extracted from greenhouse-grown leaf, stem, root, callus, and derived callus tissue. This resulted in chromosome-scale genome assemblies consisting of 10 main chromosomes as well as 126 and 165 small unplaced contigs (for BTx642 and RTx430 assemblies, respectively; **Table S1**). Both genomes are available through the Phytozome database (https://phytozome-next.jgi.doe.gov/)[29]. We then generated a synteny-guided pan-gene set consisting of the two EPICON strains in addition to the sorghum var. BTx623 reference genome (**Table S2**). This pan-gene set contained 41,607 pangenes, 25,964 of them (62.4%) shared between all three genotypes (**Fig. 2**). When comparing two genotypes, pangenes can be described as either 1:1 (one syntenic copy per genotype), 1:M (one copy in one genotype, multiple copies in the other), M:M (multiple syntenic copies in both genotypes), or 1-0/M-0 (gene uniquely present in only one genotype). While the genomes of BTx642 and RTx430 are highly similar and syntenic (**Fig. 2B,C)**, our approach identified several thousand genes that were not present in one or the other genotype, and others with copy number variation. While this could be due to inaccurate or incomplete annotations for either genome, it may represent biological differences that could underlie the phenotypic differences of these genotypes to drought (**Fig. 2A**).

**Figure 2:**
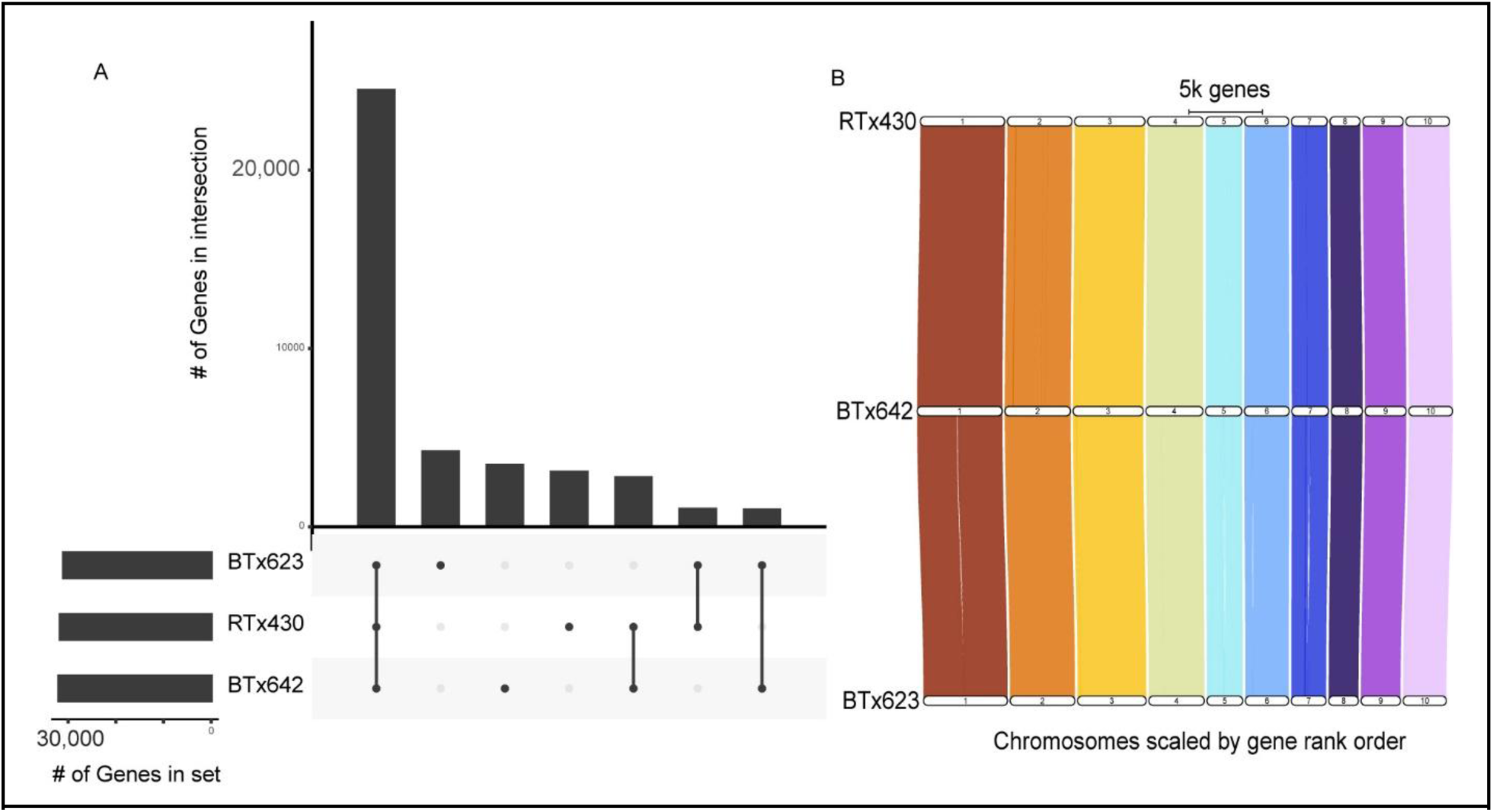
Genome similarity between three sorghum genotypes. A) UpSet plot showing numbers of genes having 1:1 relationships between the three genotypes. Dot indicates the overlap category among genes for genotypes shown on bottom left. C) Riparian plot showing high degree of synteny in gene content between three sorghum cultivars, RTx430 (top), BTx642 (middle), and BTx623 (bottom; reference).

Using this pan-gene set, we quantified transcript abundance for each sample by first mapping reads from all libraries to their corresponding reference genome. We then summarized gene expression for all pangenes (see Methods) as the sum of aligned counts for all composite genes within a pangene for each genotype. We reasoned that pangenes with multiple genes from a single genotype likely represent tandem duplication events with little functional difference. Thus, expression from any gene copy represents increased abundance in one functional gene product, making the sum of expression an appropriate summary metric. In addition to pangene expression profiles, we also compiled “genotype-centered” expression profiles for all genes in each individual genotype.

Our complete dataset consisted of 1,476 gene expression profiles of 51,865 expressed, protein-coding genes (annotated genes from either genotype, including homologs), consolidated into 23,545 expressed pangenes with representatives from both genotypes. Of these, we identified 16,358 pangenes (69.5%) that represented 1:1 orthologs. We used principal component analysis (PCA) of these 1:1 orthologs across all samples to assess the major sources of variation. We found 6 principal components representing over 85% of the variance among expressed and variable genes. These principal components apparently separated expression profiles largely by tissue, genotype, and plant age (**Fig. S1**).

### Identification of a core set of genes impacted by drought

We set out to identify genes that are differentially expressed upon drought stress consistently across years, conditions, and between genotypes, thus representing a robust drought-responsive transcriptome. To accomplish this, we used as a starting point the approach of Varoquaux *et al.* (2019), which modeled normalized expression as a set of smooth splines through sampling time (the “split-spline” approach, see Methods). We then contrasted expression models from droughted plants with those from well-watered plants. Here, we defined a gene as differentially expressed (DE) if it exhibited a FDR-controlled q-value less than 0.05 (drought vs. control across the entire timecourse in Year 2 and Year 3) and an estimated log_2_-fold change greater than 1 (drought vs. control across the time interval corresponding to pre- or post-flowering drought treatment). We repeated this analysis for each sampling year independently (“Year1”, “Year2”, “Year3”) as well as for all sampling years combined (“Year123”), aligning the time series each year by the date of anthesis (and watering transition). Through this analysis, we identified 8,503 pangenes as differentially expressed in at least one year (or all three years combined), in either genotype or drought condition, representing over 35% of the expressed pangenome. Overall, expression differences were consistent in pairwise comparisons of each year (**Fig. S2**).

Our underlying hypothesis was that, despite the meticulous care taken to execute a reproducible drought stress year-to-year and the use of plot replication in our field study, only a subset of drought DE genes would be reproducibly and differentially expressed across multiple years due to variation in growing conditions (e.g., soil effects, temperature, drought severity, etc.). Confirming our hypothesis, we observed a wide variation in the number of DE genes identified, depending on the sampling year (**Fig. 3; Fig. S3**), with only half (5,590) of the drought DE gene set being identified as DE when all three datasets are combined and analyzed together. Particularly striking was the number of genes identified as DE only in Year1. While a large number of factors may account for this phenomenon, one potential explanation is that the plot location used for Years 2-3 was different from that used in Year 1. This might result in differences in soil composition. Weather or other factors might also influence DE between years, however we did not identify a likely covariate that was consistent with our observations. For further analyses, we used DE and log-fold change estimates from a combined (Year123) analysis.

**Figure 3:**
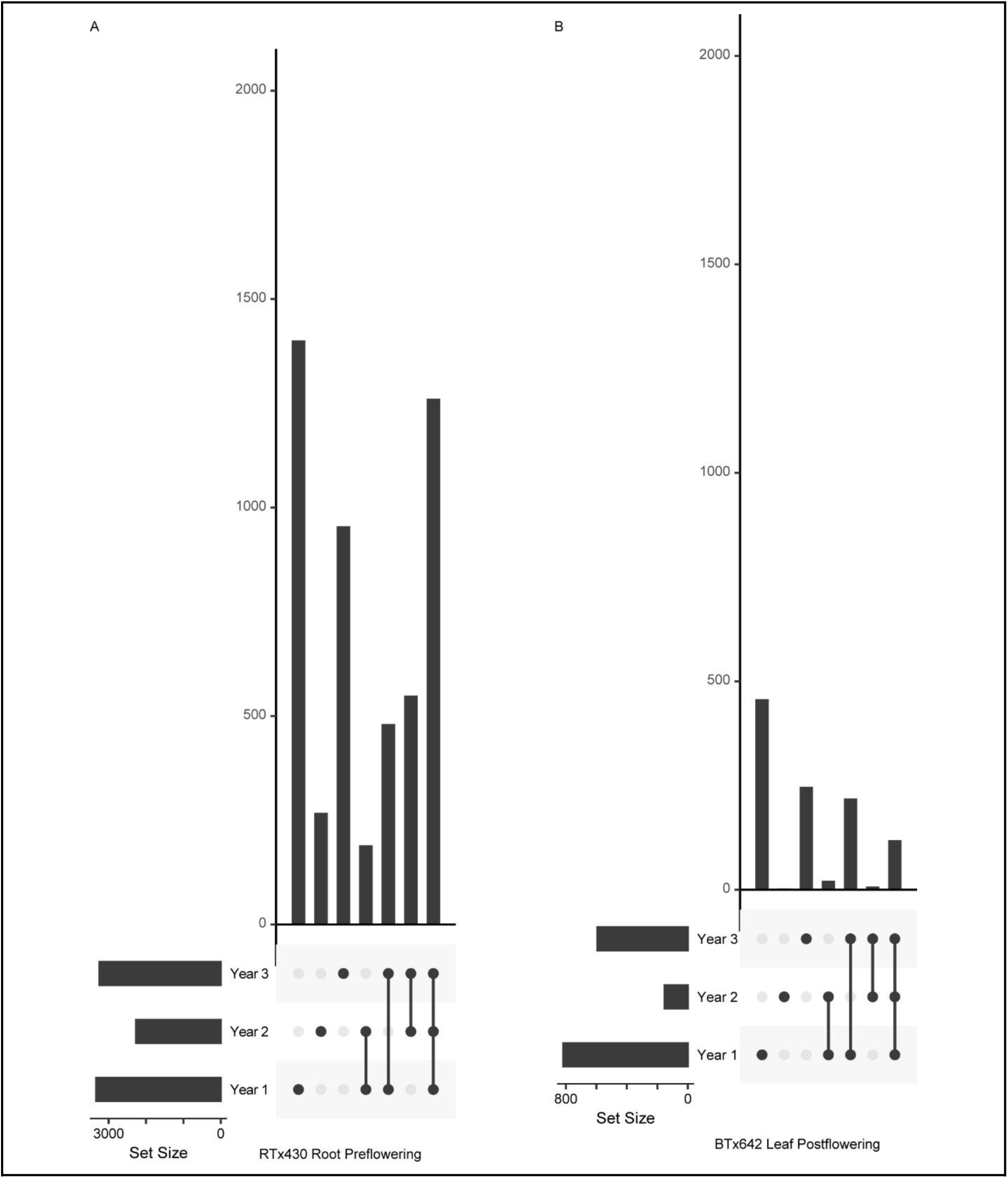
DE genes recovered from each condition per year (calculated for each year independently). Shown are selected UpSet plots describing set intersections of DE genes identified for each condition/year combination (full set of plots shown in **Fig. S3**). Individual dots represent DE genes unique to a particular year. Connected dots represent DE genes that are common to specified years.

We hypothesized that a set of genes exhibits more robust/reproducible expression changes during drought across both genotypes when considering data from all three years combined. This more heavily filtered set might enrich for genes that are less prone to variable factors, but are consistently responsive to drought. To do this, we exclusively classified genes based on whether they were differentially expressed in the same direction in both genotypes (“Core”) under Preflowering (1203 pangenes), Postflowering (703 pangenes), or during Preflowering drought recovery (“Recovery”; 935 pangenes), considering all three years of data together (though independently analyzing root and leaf data, as principal component analysis indicated this was a large source of variation in our data, **Fig. S1**). Genes assigned to the “Recovery” category did not meet the DE thresholds to classify them as drought responsive (**Fig. 4**), though we reason they are likely important for sorghum’s overall drought response. Interestingly, a comparatively small set of genes (Core “Drought” Genes, 726 pangenes, **Table S3**, **Table S4**) were found to be DE in the same direction (i.e., up- or down-regulated) in both genotypes and under both pre- and postflowering drought conditions, representing a set of universal drought-responsive genes.

**Figure 4:**
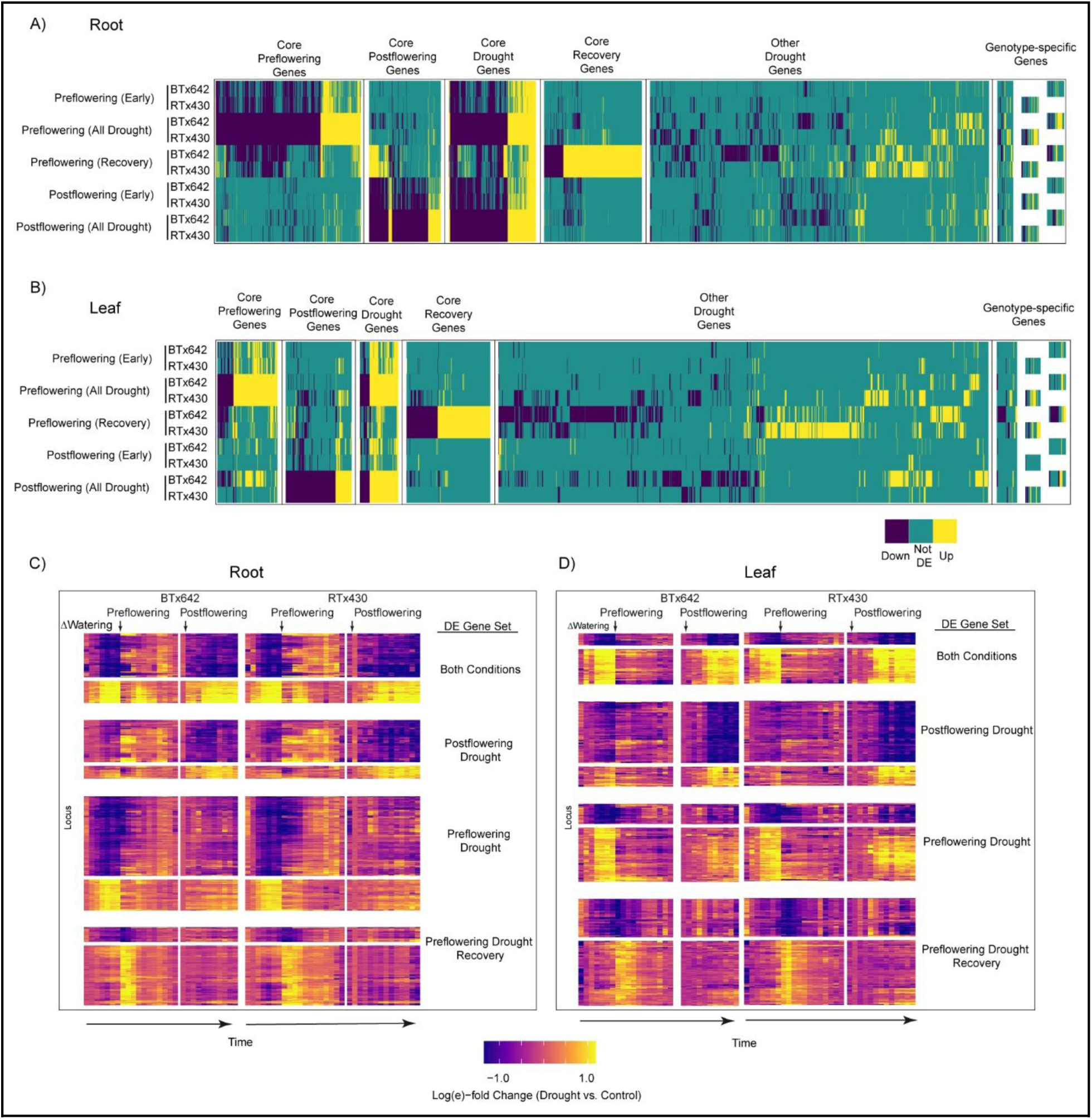
“Core” gene expression patterns. A) Genes arranged according to their categorization as Core, Other, or Genotype-specific. Core genes are either DE (in the same direction for a given tissue) in Pre-flowering, Post-flowering, Pre- and Post-flowering, or Pre-flowering Recovery in both genotypes. Other drought genes are DE but not for both genotypes (in the same direction). Genotype-specific genes are those that are expressed or present in only one genotype. These categories are assigned independently for root (a) and leaf (b) tissues. (C-D) expression profiles over the time series for Core drought genes.

We performed GO enrichment analysis on all Core genes for each drought/recovery category (**Fig. 5**). We observed significant (FDR < 0.05, BH-adjusted Fisher’s exact test) enrichment for terms associated with abiotic stress, specifically for genes up-regulated in both pre- and post-flowering drought stress. Terms related to heat, light, and ROS were represented, yet water deficiency and drought terms were notably absent among this set. Terms associated with cell wall-related processes were enriched among genes up-regulated in pre-flowering drought recovery, and commonly down-regulated in root for both pre- and post-flowering drought stress. This was also noted in a previous study [10], however this same study also measured cell wall composition and concluded that transcriptome changes among cell wall-related genes does not predict changes in actual cell wall content. We also observed enrichment of terms related to biotic stress among genes down-regulated during pre-flowering stress, consistent with our previous observation that genes associated with AMF (arbuscular mycorrhizal fungi) colonization are sharply down-regulated by drought [11]. Looking more closely at these genes, this phenomenon is recapitulated across all 3 years of this study, suggesting this is a robust feature of drought stress in sorghum roots (**Fig. S4**).

**Figure 5:**
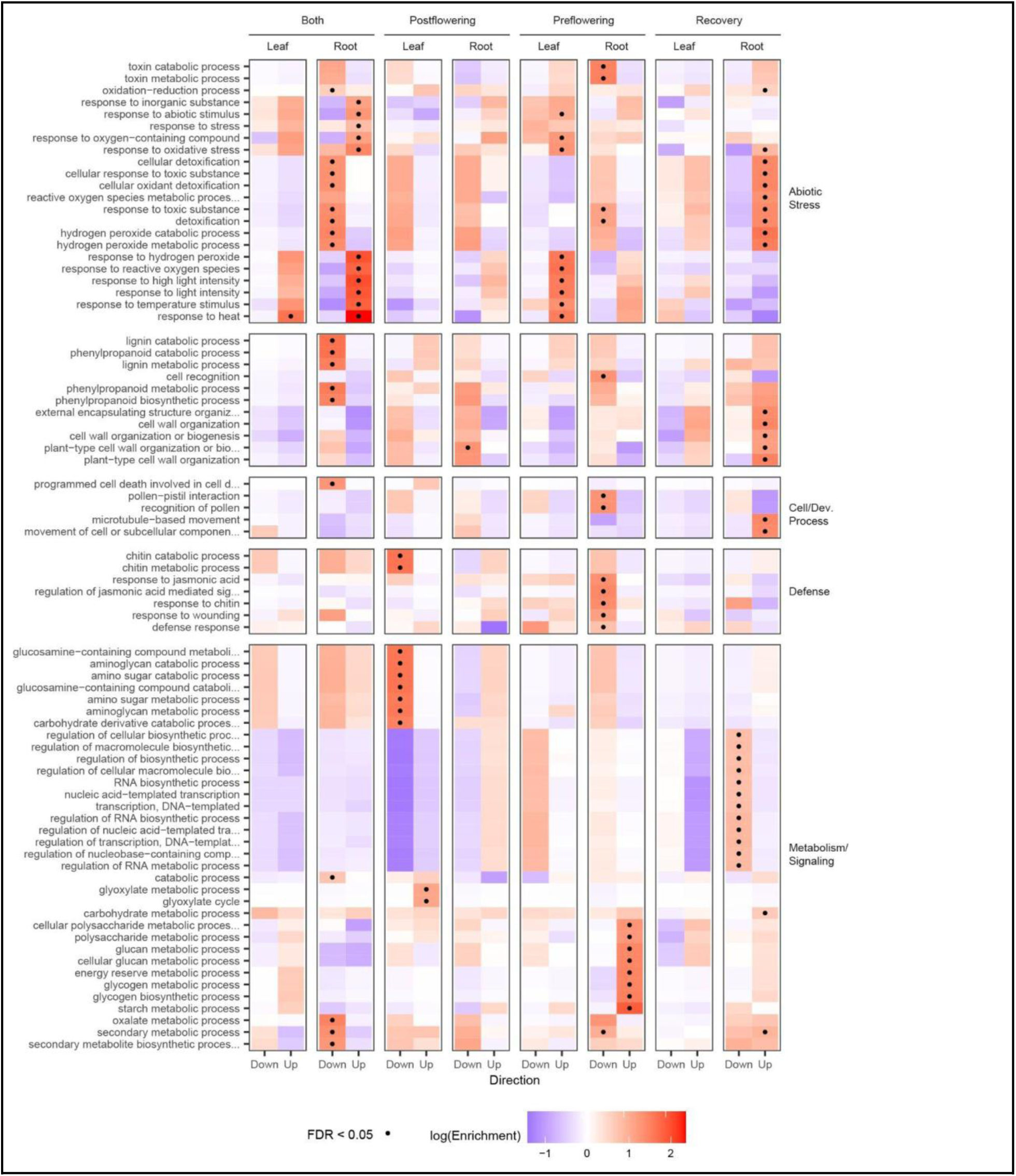
GO term enrichment for Core drought gene sets. This is broken out for genes that are up- or down-regulated in a given condition, and organized according to broad categories of GO terms. Color indicates the extent of GO term enrichment (number found significant over total number of genes with that GO term among expressed genes). Dots represent whether that enrichment is significant (Fisher’s exact test, FDR < 0.05).

To further build support for our Core gene set representing a robust set of drought-responsive genes, we turned to a recent meta-analysis [30] that also aimed to identify a set of loci in sorghum that was responsive to drought stress across multiple conditions and genotypes (including both BTx642 and RTx430). They compared drought expression data reported by 6 individual studies from sorghum grown under field or controlled conditions in different tissues at various developmental stages. They then built a model that predicts whether a gene expression profile corresponds to a drought condition and identified a set of 285 genes that are differentially expressed in sorghum, strongly contribute to that model’s accuracy, and are also conserved, differentially expressed and similarly predictive of drought in maize. We observed that a sizeable fraction (∼40%) of these genes are also differentially expressed in our study (**Fig. S7**). The Pardo predictive genes not within our set (genes strongly DE in both genotypes for a given condition) have a less extreme log-fold change distribution than those genes exclusively observed in our set under most genotype/tissue/condition combinations (adjusted pairwise Wilcoxon rank sum p-value < 0.05), but themselves are more extreme than the rest of the expressed set of sorghum genes (**Fig. S7**). That such a large number of genes appear to change expression in the same way across multiple studies lends support to these genes acting as robust markers of drought in sorghum.

Lastly, genes that did not exhibit consistent differential expression profiles across both genotypes (i.e., not “Core”) were also observed and made up a comparatively larger share of the overall number of DE genes (**Fig. 4A,B**). Other genes were identified that we designated as “genotype-specific” either by virtue of presence or absence of variation among genomes, or due to expression in one genome but not the other. These represented a small minority of the overall DE set, yet may represent interesting facets of our field study, as discussed below.

### A refined list of putative stay-green loci

One of the striking distinctions between the two genotypes in this study (and a major reason they were chosen for the EPICON project) is the ability of BTx642 to retain photosynthetic activity late into post-flowering drought stress [23,31–33]. This phenomenon has been studied for several decades, yet a comprehensive and causative list of genes underlying this phenotype, termed stay-green, has yet to be determined. Given the *de-novo* assembled genome sequence of BTx642, as well as the extensive transcriptome data generated in this study, we attempted to more specifically identify loci responsible for this agriculturally important trait. To do this, we relied on QTL mapping data generated from three studies [32–34] that are archived in the Sorghum QTL Atlas resource [35] to define rough coordinates in the BTx623 reference genome (**Fig. S6**). We then used whole genome alignment to translate these intervals to genome coordinates in our BTx642 assembly (+ a 2.5 Mbp buffer), and then identified all transcripts that fell within this region. We identified over 4,000 genes within the putative stay-green loci (**Fig. S6**). Of those genes, we found 603 that we labeled as “candidate stay-green genes”: 335 that had presence/absence variation (gene was uniquely present in BTx642 compared to RTx430), 89 that were present in both genotypes but not expressed in RTx430, and 183 that were differentially expressed between RTx430 and BTx642 (**Fig. S6**). GO term enrichment analysis of the candidate stay-green genes revealed terms related to lipid metabolism as significantly enriched, however further characterization is needed to determine whether this enrichment factors into the mechanism of the stay-green phenotype.

### Drought DE genes involved in starch and glyoxylate metabolism

We hypothesized that orthologs of Core drought genes would have evidence of drought responsiveness in model plant species. This was indeed the case, with examples including pg.02738 (Sobic.001G226600 / SbiRTX430.01G234000 / SbiBTX642.01G234500), the ortholog of a beta-amylase that accelerates starch degradation in drought stress in Arabidopsis [36]. Another instance is pg.01034 (Sobic.001G089000 / SbiRTX430.01G091100 / SbiBTX642.01G090000), the ortholog of an endoplasmic reticulum-targeted glucosidase gene critical to abscisic acid (ABA) signaling, a central drought stress pathway [37]. In roots, there was the induction of the rate-limiting step of proline biosynthesis, delta1-pyrroline-5-carboxylate synthase 1 (P5CS1, pg.16583 / Sobic.003G356000 / SbiRTX430.03G383900 / SbiBTX642.03G381100), a key osmoregulator in droughted tissue [38]. It is important to note that these examples are not exhaustive as the DE Core drought genes also include an ortholog of a gene involved in thickening of the leaf wax cuticle (pg.02691 / Sobic.001G222700 / SbiRTX430.01G230000 / SbiBTX642.01G230800), multiple heat shock proteins, and multiple dehydrins.

Some genes with well-established functions that have not previously been linked to drought responses are also present among the DE Core drought genes. One example is the induction of the glyoxylate cycle-specific enzymes, malate synthase (MLS1, pg.26913 / Sobic.006G127100 / SbiRTX430.06G133600 / SbiBTX642.06G138800) and isocitrate lyase (ICL1, pg.10765 / Sobic.002G324000 / SbiRTX430.02G328400 / SbiBTX642.02G333900) in post-flowering drought conditions (**Fig. S5**). Interestingly, the induction of ICL1 was observed in both post-flowering droughted root and leaf, whereas the induction of MLS1 was specific to post-flowering droughted leaf tissue. The function of the glyoxylate cycle is to convert acetyl-CoA, often produced via the catabolism of lipids or ketogenic amino acids, into the organic acid succinate. In fact, a key gene involved in lipid catabolism (ACX2, pg.24600 / Sobic.005G181000 / SbiRTX430.05G191100 / SbiBTX642.05G195600) shared the same DE drought pattern as the glyoxylate cycle-specific enzymes (**Fig. S5**). This observation strongly suggests that the glyoxylate cycle may have a previously unknown role in post-flowering drought tolerance.

## Discussion

### Pangenome approach

A valuable tool for plant researchers and breeders is the impressive germplasm collections for many species that contain extensive phenotypic variation resulting from local adaptations. However, recent pangenome studies have revealed extensive intraspecific presence-absence and copy number variation which complicates comparisons between even closely related lines [28,39,40]. Thus, to directly compare genomic (especially genome-wide expression) results across even closely related lines from a single species, a common gene space is needed. Traditionally, expression data from multiple genotypes have been compared to a single reference genome. While this produces acceptable results for the majority of genes for which 1 to 1 relationships exist, it under- or over-estimates expression of genes with copy number variation with respect to the reference genome and completely misses genes not found in the reference genome. Indeed, we observed that over 6,000 genes found in the two closely related sorghum genotypes (RTx430 and BTx642) used in this study do not have syntenic homologs in the sorghum reference genome produced from BTx623.

To more accurately quantify gene expression in RTx430 and BTx642, we sequenced, de-novo assembled, and annotated genomes for both of these lines. The resulting genomes were nearly complete with the only significant ambiguities contained in some centromeric regions. Indeed, most of the genome assemblies had > 50x coverage from PacBio long-read sequence data, with BUSCO scores approaching 99-100% and a relatively small number of unassembled scaffolds (< 200 for both lines). We then used a pangenome-aware approach to quantify gene expression by creating a synteny-guided pan-gene set that accounted for complex orthology relationships and presence-absence variation. Gene counts were summed across genes within each pangene/line to derive expression values. By summing read counts (instead of computing an average), we sought to account for both differential gene abundance within the genome as well as sample-to-sample variation in gene expression. This does present some caveats that we did not explore, namely that genes that are duplicated and near-identical within a genome may have suppressed read counts due to the discarding of non-uniquely mapped reads (in a 1-2 homology scenario) preferentially in one genome. We considered this a rare case, as most genes are 1:1 in these closely related genomes, and there is a greater likelihood for sequence variation among homologs in more distantly related genomes. However, this should be considered when applying a pangenome strategy to other systems. Significantly, using our pangenome approach we detected 456 genotype-specific DE genes, which underscores the importance of this approach.

### Core gene sets, major sources of variation

Using the new genomic resources, additional years of transcriptome data, and a pangenome-based strategy, we identified a robust set of genes differentially expressed in sorghum under two types of drought stress (“DE Core Genes”). This list includes a much smaller number of DE genes than many other surveys of drought in plants, primarily due to several factors. First, a stricter thresholding strategy that required genes to be DE specifically during the drought period (q < 0.05 during relevant drought intervals) with a large effect size (log2-fold change > 1) in both genotypes studied, and in the same direction. We consider this list to define a robust set of specific drought-responsive genes, rather than a comprehensive list of all genes that are potentially involved in drought response. We believe applying stringent thresholds and more narrowly defining DE categories is important, as at least for this study, a less stringent threshold for differential expression (DE between drought vs. control modeled across the entire time series with no effect size limitation) resulted in nearly all genes having some degree of differential expression, likely due to the large number of time points taken. As with many other plant species, some caution should be exercised when interpreting GO enrichment results in sorghum as a way to understand what biological processes underlie many of the DE gene sets. About 40% of the pangenes overall had some kind of GO annotation associated with them, leaving many genes (including putative stay-green loci) without putative function. This underscores a larger need for the sorghum community to experimentally ascribe functions to genes.

### Genes of interest

Using the additional two years of transcriptome data, we were able to recapitulate our previous finding with one year of data that genes associated with colonization by AMF are strongly down-regulated in pre-flowering drought. We were also able to confirm findings from our previous report [11] that major genes of the glyoxylate cycle are strongly up-regulated in post-flowering drought stress, a phenomenon that, to the best of our knowledge, has not been reported previously for any plant species. Further study will be required to determine a precise role for the glyoxylate cycle in the post-flowering drought. However, we can conjecture that mobilization of stored fatty acids and amino acids stored in leaf and root tissues via induction of the glyoxylate cycle may provide an additional source of carbon and/or nitrogen to support grain filling under conditions of drought-induced photosynthetic repression. This finding may have ramifications for understanding sorghum’s exceptional post-flowering drought tolerance, as well as potentially revealing a general post-flowering drought tolerance strategy shared by other cereal crops but overlooked in prior studies.

Lastly, we used our drought transcriptome data to further refine a list of candidate genes that may play a role in the stay-green phenotype. We did not identify a gene (or set of genes) that we can directly confirm are responsible for the phenotype. However, we did identify approximately 603 genes either differentially expressed between lines RTx430 and BTx642, or are only present in BTx642, and are also located within four broad loci that map to stay-green QTL [23,32–35] associated with the trait. These genes represent promising candidates to further characterize for their involvement in the stay-green phenotype.

### Multi-year transcriptomic datasets improve field study accuracy

A major goal of plant genetic research is to understand and improve plant traits. Yet, this often necessitates a choice between sampling plant specimens grown in tightly controlled artificial conditions in the laboratory or greenhouse or growing large numbers of plants in agricultural fields under intrinsically unpredictable conditions. Discrepancies between successful interventions under controlled conditions versus viable agricultural interventions are well understood by the community and are recognized as an important caveat to doing research in the lab [41]. Conversely, field trials often have limited predictive value due to unpredictable variability or limited sample quality or quantity. Here, we have executed a carefully controlled field study using environmental conditions that lend themselves to tight control (*e.g.,* near complete absence of summer precipitation, use of soil types well-suited for drought studies) as well as replicated field plots for two genotypes and defined drought conditions to reduce biological noise (see methods & materials). However, in spite of the planning and controls used in our study, we show that, based on the three years of transcriptomic data, conducting a field trial for a single year does not account for the substantial variation in growing conditions, including soil, weather, or other factors (e.g., plants’ biotic environment). Our observation that a basic metric (the set of genes found to be differentially regulated between drought and control conditions) is wildly different between three successive years at the same field station serves as a warning that field research, especially studies that have expression profiling as an output, should be replicated across multiple growing seasons to provide robust results.

In conclusion, we have generated one of the most comprehensive transcriptomic studies from a field-based drought trial for any plant species and by far the most comprehensive drought dataset for the drought-tolerant crop, Sorghum. By expanding our field trial through collection of samples across three growing seasons, we provide the community with a robust set of drought-responsive genes relative to the unpredictable nature of a single growing season. We also demonstrated the importance of using a pan-genome approach, especially for understanding genotype-specific traits. The dataset and our analyses are a valuable resource to the broader community that can be mined to form mechanistic hypotheses for future gene function testing and crop improvement.

## Methods

### RTx430

*S. bicolor var. RTx430 genome sequencing, assembly, and annotation.* 2500 ng of genomic DNA was sheared to >10kb using Covaris g-Tubes. The sheared DNA was treated with exonuclease to remove single-stranded ends and DNA damage repair mix followed by end repair and ligation of blunt adapters using SMRTbell Template Prep Kit 1.0 (Pacific Biosciences). The library was purified with AMPure PB beads and size selected with BluePippin (Sage Science) at >10 kb cutoff size. PacBio Sequencing primer was then annealed to the SMRTbell template library and sequencing polymerase was bound to them using Sequel Binding kit 2.0. The prepared SMRTbell template libraries were then sequenced on a Pacific Biosystems’ Sequel sequencer using v3 sequencing primer, 1M v2 SMRT cells, and Version 2.1 sequencing chemistry with 1×600 sequencing movie run times. 100 ng of DNA was sheared to 864 bp using the Covaris LE220 (Covaris) and size selected using SPRI beads (Beckman Coulter). The fragments were treated with end-repair, A-tailing, and ligation of Illumina compatible adapters (IDT, Inc) using the KAPA-Illumina library creation kit (KAPA biosystems). qPCR was used to determine the concentration of the libraries and were sequenced on the Illumina HiSeq. One Illumina 2×150 HiC library was also sequenced.

The main assembly contained 80.4x of PACBIO CCS coverage (8,091 bp average read length) that was assembled using MECAT[42] and the resulting sequence was polished using ARROW. The V4 release of *S. bicolor* var. BTx623 was broken into 58,700 nonredundant, non-overlapping 1,500 bp syntenic markers. These markers were used to identify misjoins in the RTx430 assembly. Misjoins were characterized by an abrupt change in the BTx623 linkage group. A total of 32 breaks were made to the assembly. Scaffolds were then oriented, ordered, and joined together using HIC data combined with the JUICER/JUICEBOX[43] pipeline to order and orient contigs into chromosomes. Significant telomeric sequence was properly oriented in the assembly. A total of 553 joins were applied to the broken assembly to form the final assembly consisting of 10 chromosomes. 97.9% of the assembled sequence is contained in the chromosomes. Additionally, homozygous SNPs and INDELs were corrected in the release sequence using ∼60x of Illumina reads (2×150, 500bp insert).

Transcript assemblies were made from ∼5.4B pairs of 2×150 stranded paired-end Illumina RNA-seq reads using PERTRAN (Shu, unpublished). About 1.5M PacBio Iso-Seq CCSs were corrected and collapsed by genome guided correction pipleine (Shu, unpublished) to obtain ∼292K putative full-length transcripts. 192,495 transcript assemblies were constructed using PASA[44] from RNA-seq transcript assemblies, CCSs above and ESTs from NCBI. Loci were determined by transcript assembly alignments and/or EXONERATE[45] alignments of proteins from *Arabidopsis thaliana*, soybean, rice, *Setaria viridis*, grape, high confidence and transcriptome supported BTx623 gene models and Swiss-Prot proteomes to repeat-soft-masked RTx430 genome using RepeatMasker[46] with up to 2K BP extension on both ends unless extending into another locus on the same strand. Gene models were predicted by homology-based predictors, FGENESH+[47], FGENESH_EST (similar to FGENESH+, but using EST to compute splice site and intron input instead of protein/translated ORF), and EXONERATE[45], PASA assembly ORFs (in-house homology constrained ORF finder) and from AUGUSTUS via BRAKER1[48]. The best scored predictions for each locus are selected using multiple positive factors including EST and protein support, and one negative factor: overlap with repeats. The selected gene predictions were improved by PASA[44]. Improvement includes adding UTRs, splicing correction, and adding alternative transcripts. PASA-improved gene model proteins were subject to protein homology analysis to above mentioned proteomes to obtain Cscore and protein coverage. Cscore is a protein BLASTP score ratio to MBH (mutual best hit) BLASTP score and protein coverage is highest percentage of protein aligned to the best of homologs. PASA-improved transcripts were selected based on Cscore, protein coverage, EST coverage, and its CDS overlapping with repeats. The transcripts were selected if its Cscore is larger than or equal to 0.5 and protein coverage larger than or equal to 0.5, or it has EST coverage, but its CDS overlapping with repeats is less than 20%. For gene models whose CDS overlaps with repeats for more than 20%, their Cscore must be at least 0.9 and homology coverage at least 70% to be selected. The selected gene models were subject to Pfam analysis and gene models whose protein is more than 30% in Pfam TE domains were removed and weak gene models.

Incomplete gene models, low homology supported without fully transcriptome supported gene models and short single exon (< 300 BP CDS) without protein domain nor good expression gene models were manually filtered out.

## BTx642

*S. bicolor var. BTx642 genome sequencing, assembly, and annotation.* 2500 ng of genomic DNA was sequenced as described for RTx430. The main assembly consisted of 52.36x of PACBIO coverage (7,024 bp average read size), and was assembled using MECAT[42] and the resulting sequence was polished using ARROW. A total of 61,546 unique, non-repetitive, non-overlapping 1 KB syntenic markers were generated using the version 3.0 *S. bicolor* genome release and aligned to the polished BTx642 assembly. Contig breaks were identified as an abrupt change in linkage group. A total of 16 breaks were made. The broken contigs were then ordered, oriented, and assembled into 10 chromosomes using the version 3.0 S. bicolor syntenic markers. A total of 155 joins were made during this process. The HiC library was aligned to the integrated chromosomes, and several minor rearrangements were made. Adjacent alternative haplotypes were identified on the joined contig set. Althap regions were collapsed using the longest common substring between the two haplotypes. A total of 36 adjacent altHaps were collapsed. Care was taken to ensure that telomere was properly oriented in the chromosomes, and the resulting sequence was screened for retained vector and/or contaminants. Finally, Homozygous SNPs and INDELs were corrected in the release sequence using ∼55.4x of Illumina reads (2×150, 400bp insert).

Transcript assemblies were made from ∼349M pairs of 2×150 stranded paired-end Illumina RNA-seq reads and ∼2B pairs of 2×150 stranded paired-end Illumina RNA-seq reads (expression profile experiments) using PERTRAN (Shu, unpublished). About 1.5M PacBio Iso-Seq CCSs were corrected and collapsed by genome guided correction pipleine (Shu, unpublished) to obtain ∼1M putative full-length transcripts. 215,476 transcript assemblies were constructed using PASA[44] from RNA-seq transcript assemblies above. Loci were determined by transcript assembly alignments and/or EXONERATE[45] alignments of proteins from *Arabidopsis thaliana*, soybean, rice, Setaria viridis, aquillegia, grape and Swiss-Prot proteomes to repeat-soft-masked BTx642 genome using RepeatMasker[46] with up to 2K BP extension on both ends unless extending into another locus on the same strand. Gene models were predicted by homology-based predictors, FGENESH+[47], FGENESH_EST (similar to FGENESH+, but using EST to compute splice site and intron input instead of protein/translated ORF), and EXONERATE[45], PASA assembly ORFs (in-house homology constrained ORF finder) and from AUGUSTUS via BRAKER1[48]. The best scored predictions for each locus are selected using multiple positive factors including EST and protein support, and one negative factor: overlap with repeats. The selected gene predictions were improved by PASA. Improvement includes adding UTRs, splicing correction, and adding alternative transcripts. PASA-improved gene model proteins were subject to protein homology analysis to above mentioned proteomes to obtain Cscore and protein coverage. Cscore is a protein BLASTP score ratio to MBH (mutual best hit) BLASTP score and protein coverage is highest percentage of protein aligned to the best of homologs. PASA-improved transcripts were selected based on Cscore, protein coverage, EST coverage, and its CDS overlapping with repeats. The transcripts were selected if its Cscore is larger than or equal to 0.5 and protein coverage larger than or equal to 0.5, or it has EST coverage, but its CDS overlapping with repeats is less than 20%. For gene models whose CDS overlaps with repeats for more than 20%, their Cscore must be at least 0.9 and homology coverage at least 70% to be selected. The selected gene models were subject to Pfam analysis and gene models whose protein is more than 30% in Pfam TE domains were removed and weak gene models. Incomplete gene models, low homology supported without fully transcriptome supported gene models and short single exon (< 300 BP CDS) without protein domain nor good expression gene models were manually filtered out.

### DNA and Iso-seq

RNA for Iso-seq was prepared from mature root, leaf, panicle, stem and callus tissues from RTx430 and BTx642 plants grown in greenhouse conditions. Briefly, tissue was flash-frozen in liquid nitrogen and kept at -80 deg. C or on dry ice until RNA extraction. Prior to extraction, 1-2 g of tissue was plunged in liquid nitrogen, then ground in a Retsch bead mill (Tissuelyser) using 35 mL grinding jar with 2×12mm stainless steel grinding balls for 5 minutes at 30 Hz. Jars were re-frozen in liquid nitrogen, rotated, then ground again for an additional 5 minutes at 30 Hz.

Tissue was scraped out of the jars, and 500 mg was measured into 2 mL tubes. RNA was extracted from each tissue aliquot using the Qiagen RNeasy kit (Cat. #AM217004) according to the manufacturer’s specifications. RNA was submitted to the Joint Genome Institute for Iso-seq library creation and sequencing. High molecular weight DNA was extracted at Arizona Genomics Institute.

### Growth conditions and drought treatment

Sorghum plants were planted, grown, and treated as described by Varoquaux et al (2019) [11] with some differences in years 2 and 3. These included, for year 2, plants were sown in pre-wetted plots at Kearny Agricultural Research and Extension Center (Parlier, CA) on June 6th, 2017, and Week 1 samples were collected on June 28th, 2017, 16 days after emergence).

Additional samples were acquired 8 hours, 1 and 3 days after watering resumed (August 11th, 2027) from control and pre-flowering drought-treated plots, as well as 2 and 5 days after water cessation (August 11th, 2017) from control and post-flowering plots. Otherwise, sampling dates occurred every 7 days until the conclusion of the experiment on October 11th, 2017. Year 3 plants were grown similar to year 2, with seeds sown on June 8th, 2018, Week 1 samples collected on June 26th, and the water transition occurring on August 15th, 2018.

### Sample collection, RNA isolation, library sequencing, and quantification

All samples were collected and processed as described [11]. RNA was extracted as described [11]. Sequencing was performed at the Joint Genome Institute (Award DOI: 10.46936/10.25585/60001015), targeting 80M reads per library. Raw FASTQ file reads were filtered and trimmed using the JGI QC pipeline resulting in the filtered FASTQ file. Using BBDuk[49], raw reads were evaluated for artifact sequences by kmer matching (kmer=25), allowing 1 mismatch. Detected artifacts were trimmed from the 3’ end of the reads. RNA spike-in reads, PhiX reads and reads containing any Ns were removed. Quality trimming was performed using the phred trimming method set at Q6. Following trimming, reads under the length threshold were removed (1/3 of the original read length). Filtered reads from each library were aligned to the RTx430 or BTx642 reference genome using HISAT2 version 2.1.0[50]. featureCounts[51] was used to generate the raw gene counts file using gff3 annotations. Only primary hits assigned to the reverse strand were included in the raw gene counts (-s 2 -p -- primary options).

### Pangenome construction and summary

The GENESPACE R package[52] (version 0.8.5) was used to generate a pangenome from three sorghum genomes: *S. bicolor* v3.1.1[26], RTx430 v. 2.1 and BTx642 v.1.1 using default settings. Gene counts from genes within pangenes were summed to generate pangene expression profiles. Pangenes that lacked representation from both RTx430 and BTx642 were excluded from additional analyses.

### Gene and pangene summary statistics and differential expression testing

The Moanin R package ([53]) was used to identify differentially expressed genes using several different contrasts. First, DE was tested for drought vs. control in either genotype or tissue using all time points (overall DE). Second, DE was tested using a subset of time points corresponding to pre-flowering drought (prior to 57 DAG) or post-flowering drought (after 63 DAG). Summary statistics, including an estimated log2-fold change and a corresponding adjusted (BH) p-value were computed for each test. DE genes for a particular contrast were identified as being significantly different between drought vs. control (overall DE p < 0.01) and had a log2-fold change (absolute value) greater than 1. This was computed for pangenes (using expression for syntenic, homologous genes in both genotypes) and using genotype-specific expression data. GO enrichment was performed using the topGO R package [54] using all expressed genes (at least 5 libraries with at least 1 count) or pangenes (more than 1% of libraries with at least 1 count) as background.

### Genome alignment and STG locus identification

The BTx642, BTx623 and RTx430 genomes were aligned using the minimap2[55] tool using default parameters associated with the “asm5” preset. Stay-green coordinates were first identified for BTx623 using the Sorghum QTL Atlas[35] for the top 4 loci on chromosomes 2, 3 and 5. These coordinates were then mapped to the BTx642 genome using the whole-genome alignment, filtering for hits that had a minimum alignment quality equal to 60 on the positive strand, adding a 2.5 Mbp extension on either side. Genes found within these intervals on the BTx642 genome were identified as potential stay-green candidates for further analysis.

## Data availability

Raw sequencing data is available at the National Institute of Biotechnology Information Sequence Read Archive (NCBI SRA). SRA Accessions for each sample are available in Supplementary Table S5. Processed data matrices as well as analysis code is available at https://code.jgi.doe.gov/BJCole/epicon-3-year-transcriptome.

## Acknowledgements

This project was supported in part by DOE Grant DE-SC0014081 awarded to P.G.L., D.C.D., J.D., R.H., E.P., A.V., and J.V.. This project was also supported by an Early Career Research Program award to B.C. The work (proposal: 10.46936/10.25585/60001015) conducted by the U.S. Department of Energy Joint Genome Institute (https://ror.org/04xm1d337), a DOE Office of Science User Facility, is supported by the Office of Science of the U.S. Department of Energy operated under Contract No. DE-AC02-05CH11231

---

**Figure S1:**
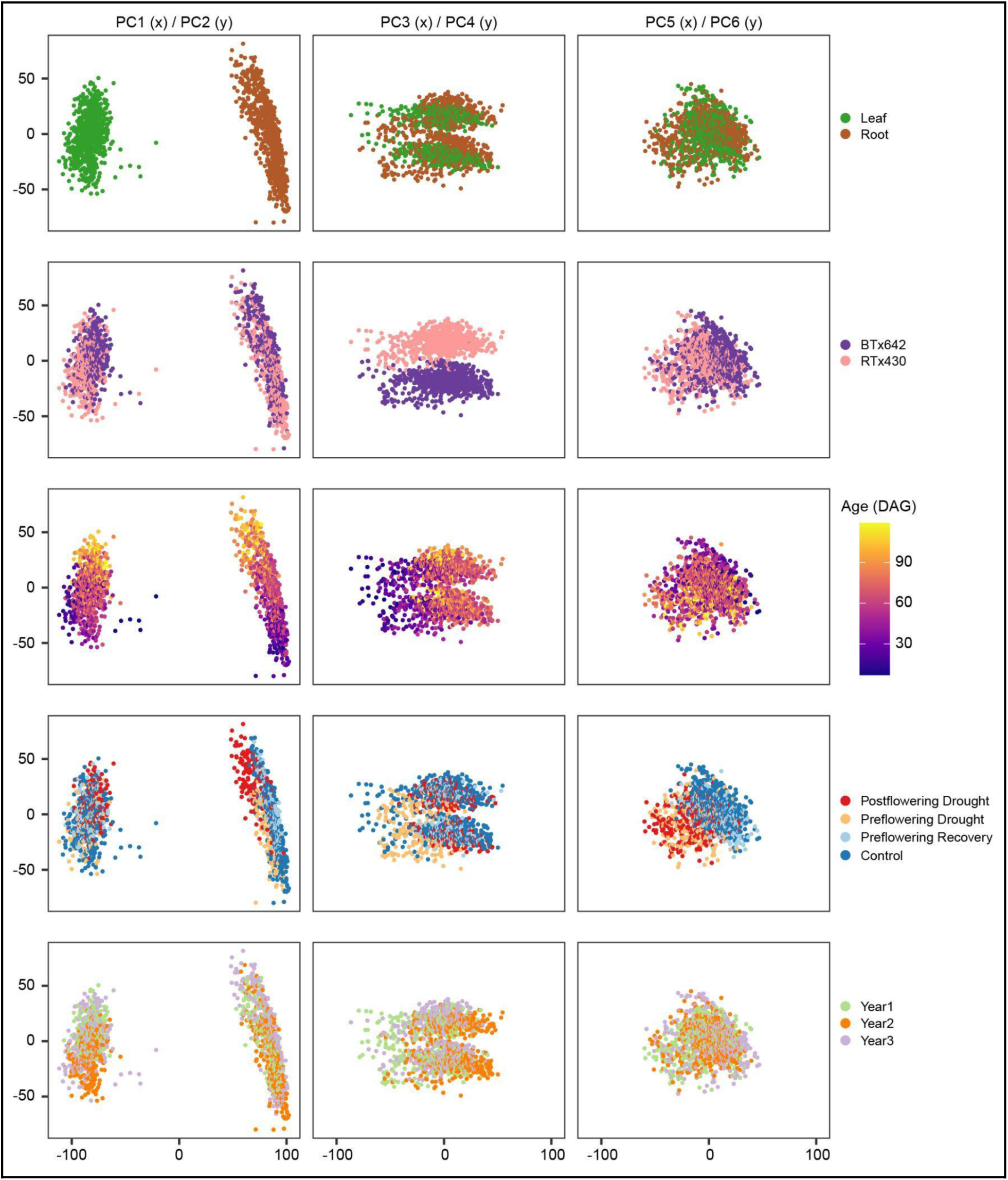
Variation between datasets. PCA was conducted with all transcriptomes, with the first six principal components plotted.

**Figure S2:**
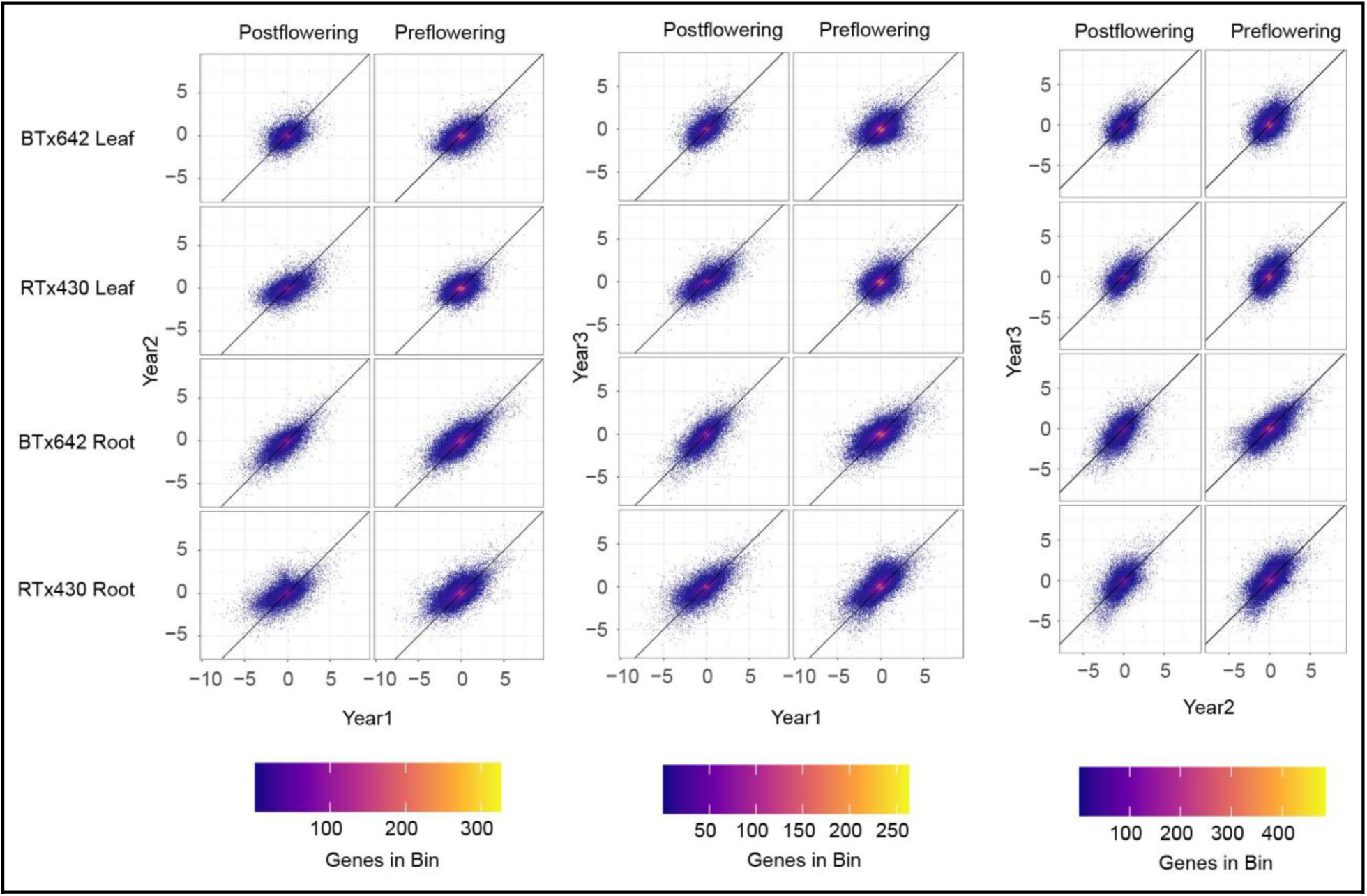
Pairwise comparison of log-fold change in gene expression by year. Shown are log2-fold changes of each gene in pairwise comparisons across years. Left, Year1 vs. Year2; middle, Year1 vs. Year3; Right, Year2 vs. Year3. Color scale represents point density (2-dimensional binning).

**Figure S3:**
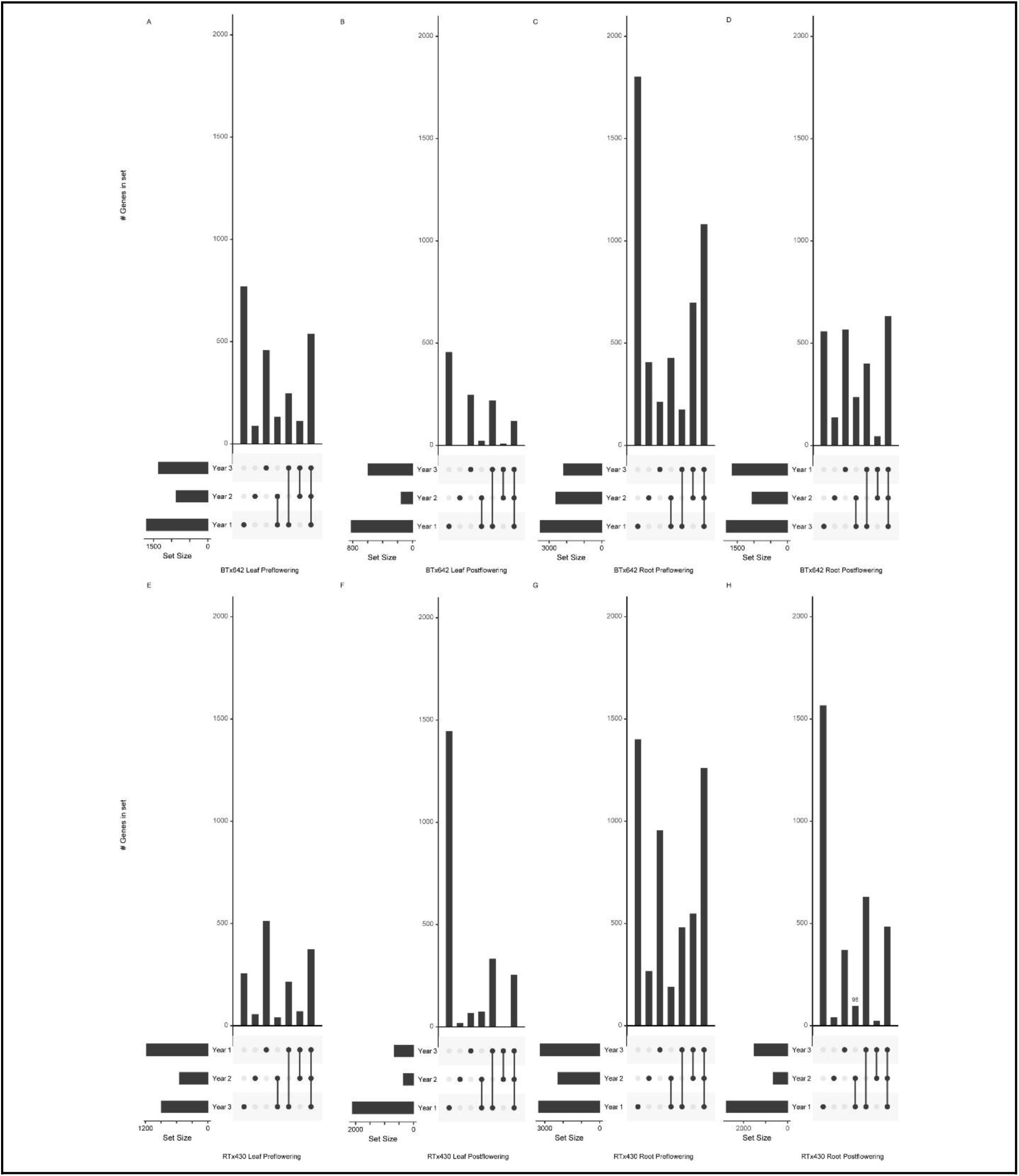
DE genes recovered from each condition per year. Shown are UpSet plots describing set intersections of DE genes identified for each condition/year combination. Individual dots represent DE genes unique to a particular year. Connected dots represent DE genes that are common to specified years. This is an expanded set of comparisons, similar to Fig. 3.

**Figure S4:**
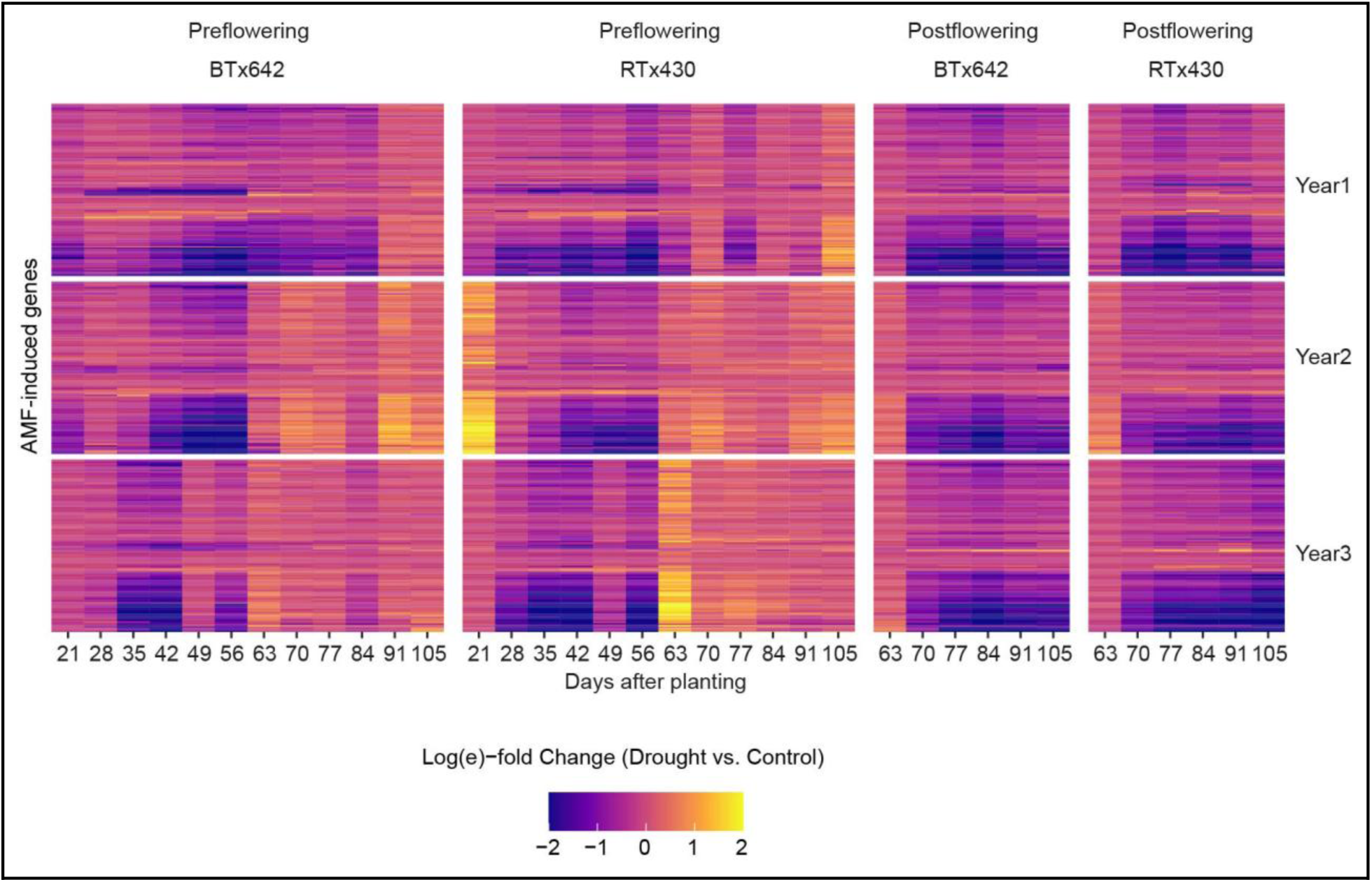
Root expression of AMF-related genes decreases during drought. Genes known to be induced by AMF are sharply down-regulated during both pre- and post-flowering drought stress in sorghum roots. Shown is a heat map of log-fold change expression across a time course for all three sampling years.

**Figure S5:**
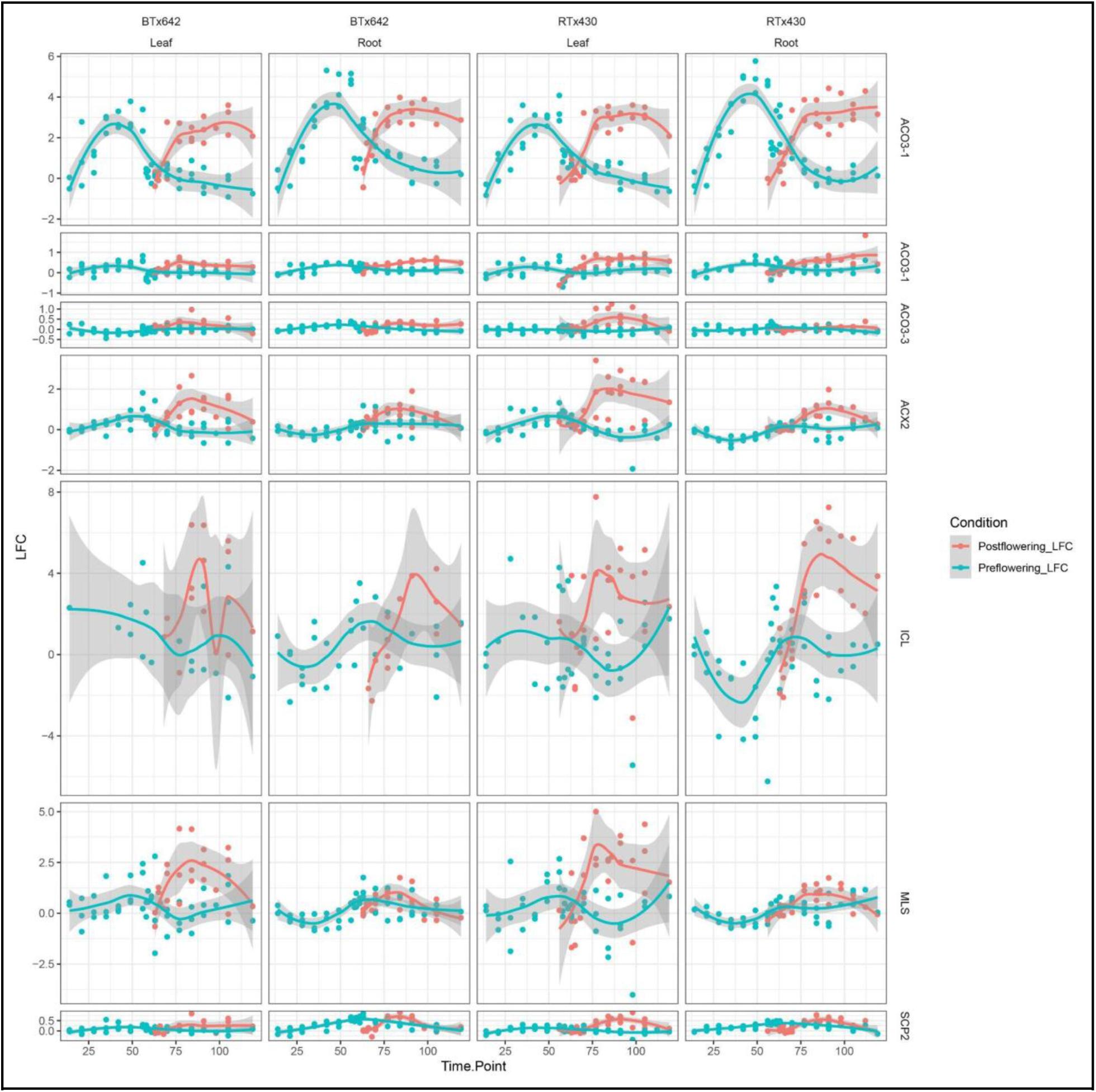
Expression of 7 pangenes corresponding to glyoxylate pathway loci in sorghum. Most genes are up-regulated in post-flowering drought stress, though several are also up-regulated during pre-flowering drought.

**Figure S6:**
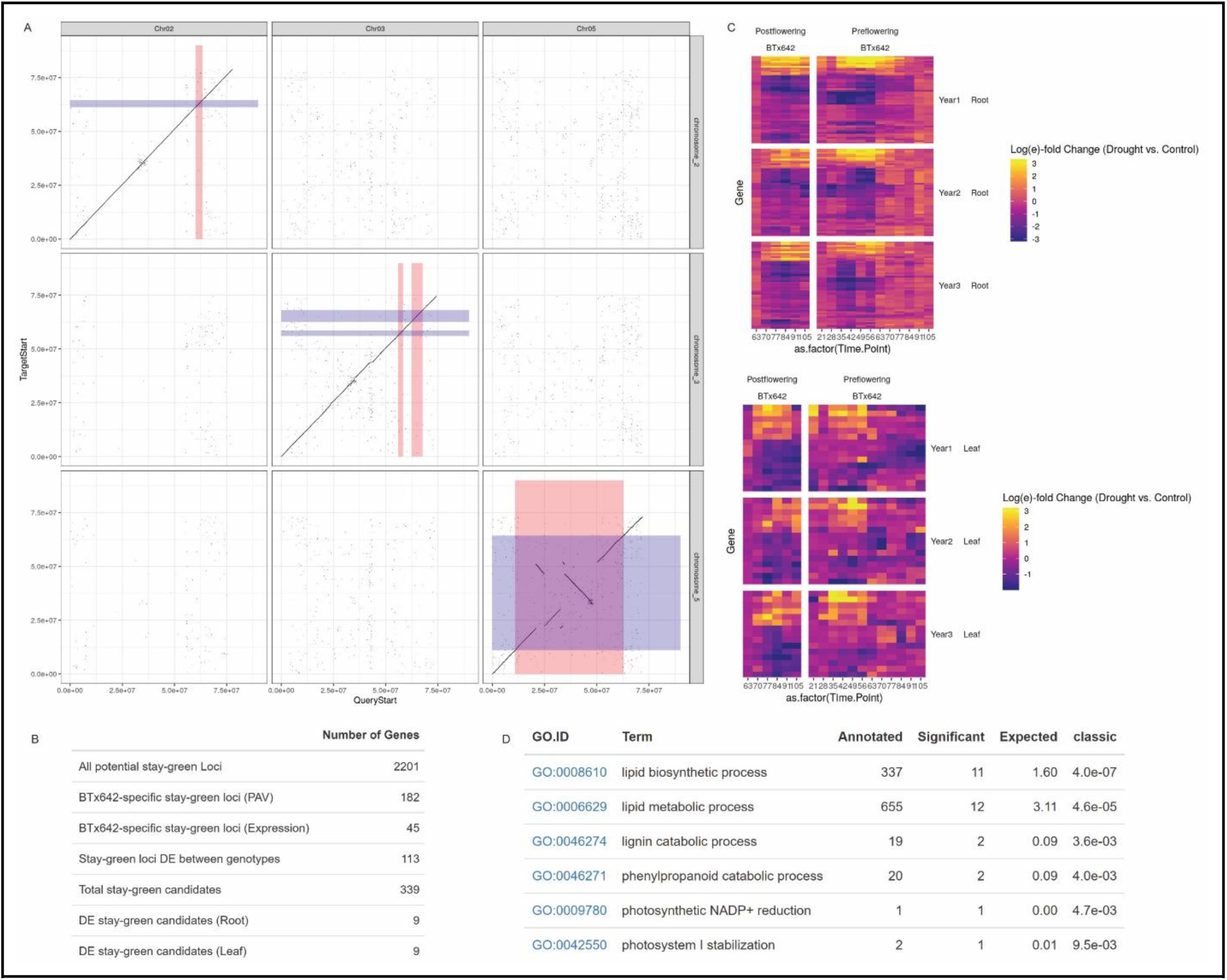
Identification of stay-green candidates in EPICON data. Potential stay-green loci were first identified by comparing the BTx623 genome to BTx642, then using these alignments to map coordinates of 4 major QTL onto the BTx642 genome (A). Genes existing within these coordinates were then further characterized as having presence-absence variation (PAV) between the RTx430 and BTx642 genomes, being exclusively expressed in the BTx642 genome, being DE between genomes, and being DE in drought in either root or leaf in BTx642. The “Total stay-green candidates” were the union set of those Stay-Green associated genes that had PAV or specific expression, or that were significantly DE between genotypes (B). Expression profiles for these genes are plotted in (C) for those DE in Root or Leaf. GO term enrichment was also performed for these genes (D).

**Figure S7:**
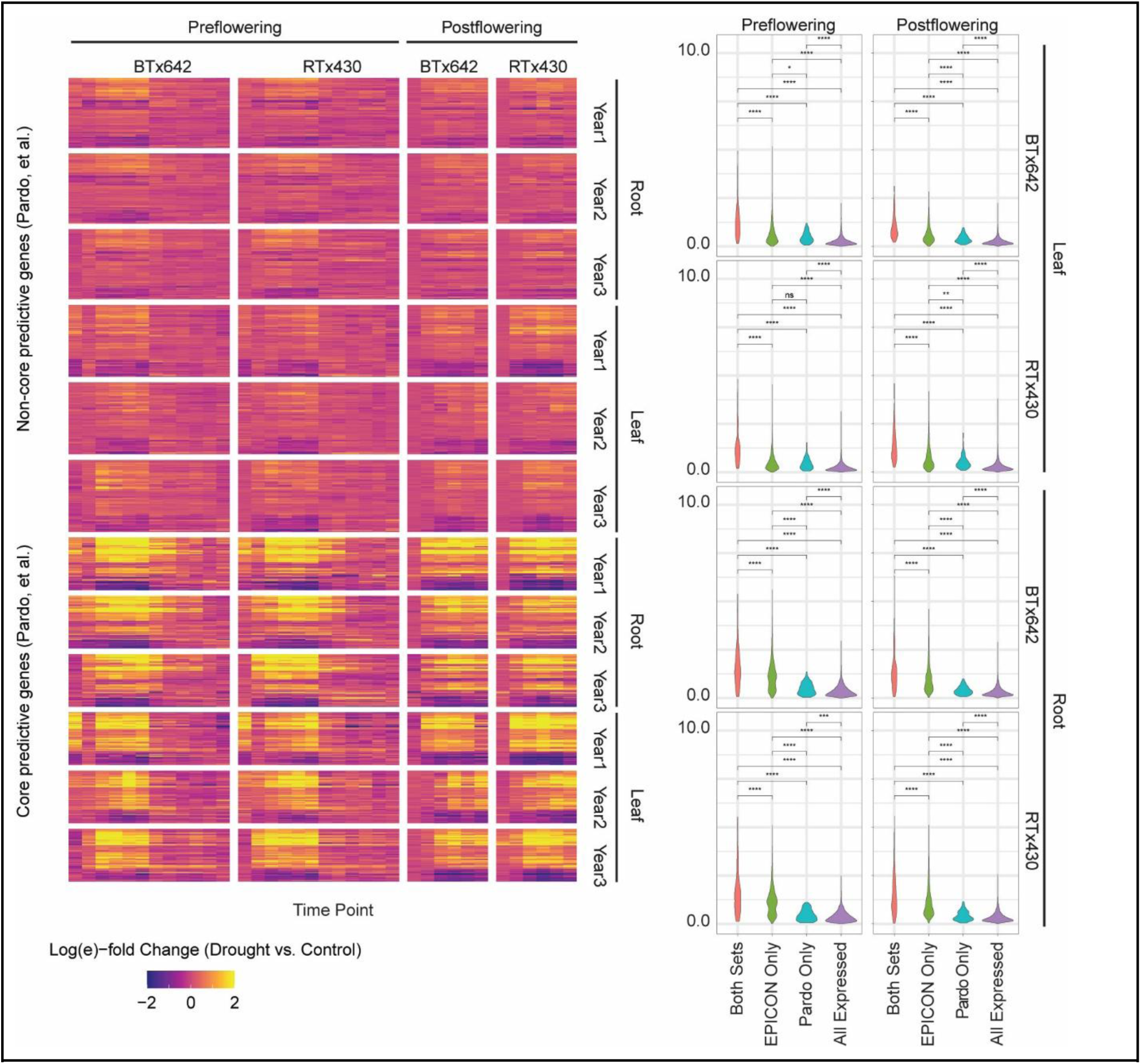
Comparison to meta-analysis of sorghum drought transcriptomes. Core drought genes were compared to genes identified as important for drought (Pardo, et al., 2023). The left panel shows a heatmap of expression of genes identified by Pardo, et al. that were either identified or not as Core drought genes in the present study. The right panel shows a violin plot of expression for the Core drought and Pardo predictive gene sets, compared to the set of all expressed genes. Asterisks indicate significance levels for pairwise Wilcoxon Rank Sum tests (p < 0.05*; p <

